# A CRISPR screen identifies redox vulnerabilities for KEAP1/NRF2 mutant non-small cell lung cancer

**DOI:** 10.1101/2022.03.08.483452

**Authors:** Chang Jiang, Nathan P. Ward, Nicolas Prieto-Farigua, Yun Pyo Kang, Anish Thalakola, Mingxiang Teng, Gina M. DeNicola

## Abstract

The redox regulator NRF2 is hyperactivated in a large percentage of non-small cell lung cancer (NSCLC) cases, which is associated with chemotherapy and radiation resistance. To identify redox vulnerabilities for KEAP1/NRF2 mutant NSCLC, we conducted a CRISPR-Cas9-based negative selection screen for antioxidant enzyme genes whose loss sensitized cells to sub-lethal concentrations of the superoxide (O_2_•^−^)-generating drug β-Lapachone. While our screen identified expected hits in the pentose phosphate pathway, the thioredoxin-dependent antioxidant system, and glutathione reductase, we also identified the mitochondrial superoxide dismutase 2 (SOD2) as one of the top hits. Surprisingly, β-Lapachone did not generate mitochondrial O_2_•^−^ but rather SOD2 loss enhanced the efficacy of β-Lapachone due to loss of iron-sulfur protein function, loss of mitochondrial ATP maintenance and deficient NADPH production. Importantly, inhibition of mitochondrial electron transport activity sensitized to β-Lapachone, demonstrating these effects may be translated to increase ROS sensitivity therapeutically.

## INTRODUCTION

Lung cancer is the second most common cancer and the leading cause of cancer death worldwide. Non-small-cell lung cancer (NSCLC) accounts for 80-85% of all diagnosed lung cancer cases [1], of which a third have mutations in the NRF2/KEAP1 circuit [2]. NRF2 (Nuclear factor erythroid 2 p45-related factor or NFE2L2) is a stress-responsive transcription factor that regulates the response to oxidative stress. KEAP1 (Kelch-like ECH-associated protein) directs NRF2 degradation, and accordingly, disruption of KEAP1 increases the abundance and activity of NRF2. Aberrant NRF2 activation promotes cancer progression, metastasis and confers profound resistance to chemo and radiotherapy [1, 3–11]. Thus, targeting NRF2 and its downstream processes holds great promise for lung cancer therapy development. However, no effective treatment strategy targeting NRF2/KEAP1 has been developed for clinical use. Direct therapeutic targeting of NRF2 is challenging due to its lack of catalytic domains. High-throughput drug screens have been performed to identify NRF2 inhibitors, but they lacked selectivity or demonstrated systemic toxicity [12, 13]. It is therefore critical to identify and characterize vulnerabilities of KEAP1/NRF2 mutant NSCLC to develop effective therapies for patients harboring these mutations.

NRF2 controls the transcription of a plethora of antioxidant enzymes, thereby regulating the detoxification of reactive oxygen species (ROS) [4, 14, 15]. These enzymes include the glutathione (GSH)- and thioredoxin (TXN)-dependent antioxidant systems, heme and ion metabolic enzymes, xenobiotic detoxification enzymes, and enzymes that produce the reduced form of nicotinamide adenine dinucleotide phosphate (NADPH). NADPH is the major cellular reducing power that supports both antioxidant defenses and reductive biosynthesis. The pentose phosphate pathway (PPP) is the largest contributor to cytosolic NADPH in most cultured mammalian cells [16–18]. NRF2 activation promotes the expression of PPP enzymes, including glucose-6-phsphate dehydrogenase (G6PD) and 6- phosphogluconate dehydrogenase (PGD), transaldolase and transketolase (TKT) [19], thereby maintaining high NADPH levels. PPP enzymes support NRF2-induced proliferation [19–21] and CRISPR screening revealed the dependence of NRF2 hyperactive cancer cells on the PPP for survival [22, 23]. Moreover, NRF2 activation impacts parallel, redundant antioxidant systems that function in multiple intracellular compartments. These thiol-dependent antioxidant pathways are mediated by the antioxidants GSH and TXN, and their respective reductases, namely GSH reductase (GSR) and thioredoxin reductases (TXNRD1 and TXNRD2), which both require NADPH as their reducing agent. The GSH/GSR and TXN/TXNRD pathways critically regulate and maintain cellular thiol redox homeostasis and protein dithiol/disulfide balance [24]. Indeed, we previously reported that NSCLC cell lines harboring NRF2/KEAP1 mutations demonstrated elevated resistance to the superoxide-generating small molecule β-Lapachone [25]. Inhibition of the thioredoxin-dependent antioxidant system or SOD1 could overcome NRF2-mediated resistance to β-Lapachone, but inhibition of glutathione synthesis had no effect [25]. However, which individual antioxidant enzymes mediate β-Lapachone resistance downstream of NRF2 is unknown. Other studies using KEAP1/NRF2 wild-type cells have shown that inhibition of Peroxiredoxin 1, Peroxiredoxin V or MTHFD2 can increase β-Lapachone cytotoxicity [26–28], suggesting that antioxidant enzymes can play non-redundant functions in ROS detoxification. Due to the redundancy and complexity of NRF2 induced ROS detoxification program, a systematic study strategy is critically needed to identify key vulnerabilities that contribute to the resistance of NSCLC to ROS. To this end, we conducted CRISPR-based negative screens to identify antioxidant genes whose loss exacerbate the β-Lapachone induced cell death in NSCLC with NRF2 hyperactivation.

## 2. RESULTS

### 2.1 CRISPR/Cas9 screens identify known and novel vulnerabilities of NSCLC cells to oxidative stress induced by β-Lapachone

We previously reported that NRF2 activation in NSCLC confers profound resistance to ROS [25]. To discover synthetic lethal target genes whose loss exacerbates the ROS-induced cell death in NSCLC with NRF2 hyperactivation, we conducted CRISPR-Cas9-based negative selection screens for antioxidant enzyme genes whose loss sensitized NSCLC cells to sub-lethal concentrations of β-Lapachone (Supplementary Fig 1A). Human KEAP1 mutant NSCLC cell lines A549 and HCC15 were transduced with a single guide RNA (sgRNA) library targeting 139 antioxidant enzyme genes containing ~12 unique sgRNAs per gene, and treated with either vehicle control (DMSO) or β-Lapachone. To recapitulate β-Lapachone *in vivo* half-life conditions [29], cells were cultured with either DMSO or β-Lapachone for 2 h every other day for 3 sequential treatments. At the end of the culture period, deep sequencing was used to measure the abundance of all sgRNAs in the DMSO and β-Lapachone-treated cells (Fig. 1A). In parallel to the negative selection screen, we also conducted positive-selection CRISPR-based screens on A549 and HCC15 cells for antioxidant enzymes whose loss allowed cell survival under treatment of higher concentrations of β-Lapachone (Fig. S1A). Importantly, NQO1, which is essential for ROS generation by β-Lapachone and whose loss promotes resistance to β-Lapachone (Fig. S1B), was identified as the top hit in both A549 and HCC15 positive-selection screens (Fig. S1C), supporting the rigor of the screen.

**Fig. 1.**
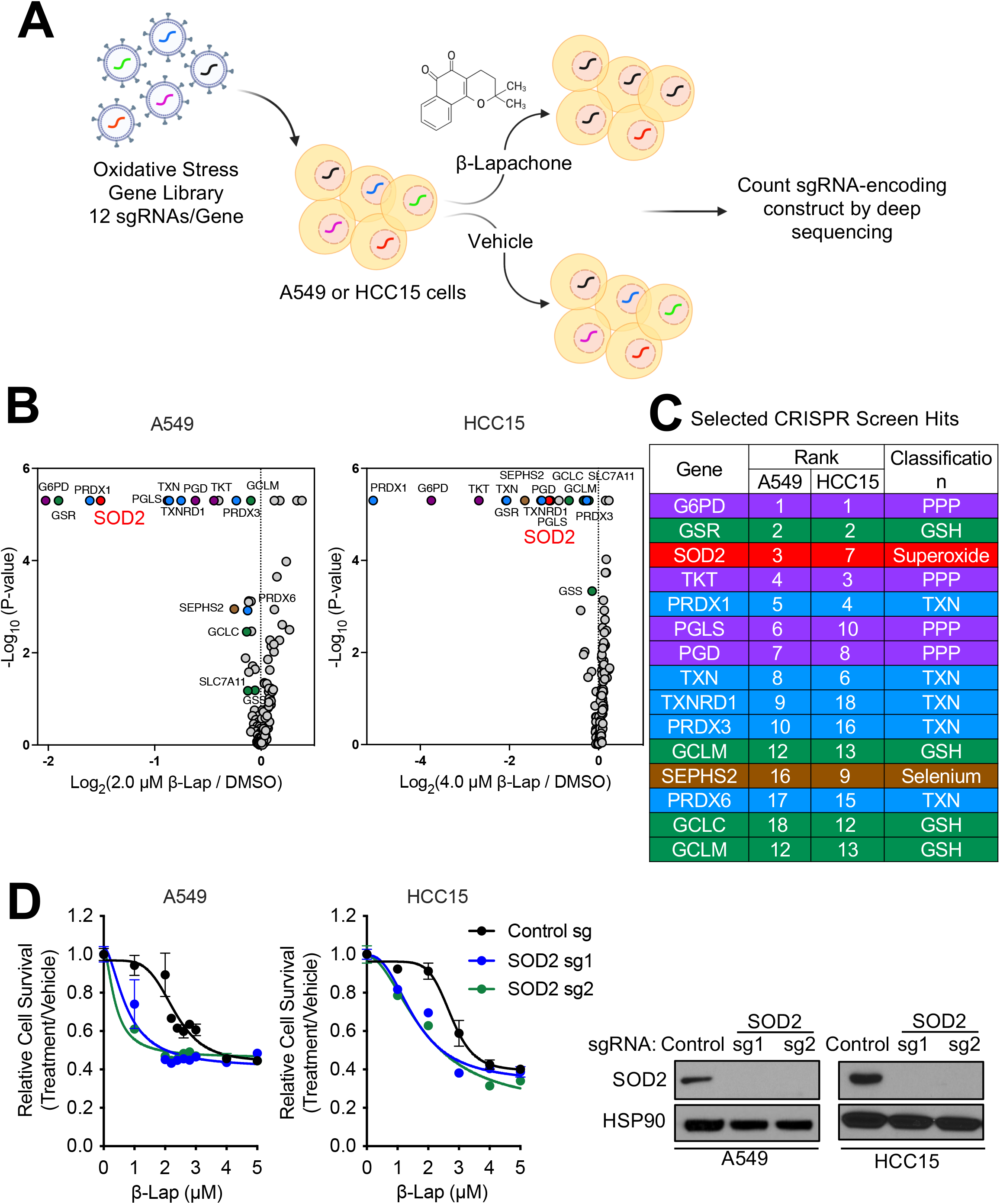
CRISPR/Cas9 screens identify redox vulnerabilities of KEAP1 mutant NSCLC cells. (A) Schematic of the antioxidant enzyme gene focused pooled sgRNA library screen. KEAP1^MUT^ A549 and HCC15 cells were infected with sgRNA library lentivirus at a MOI of ~ 0.3. After selection with puromycin, cells were treated with either vehicle control (DMSO) or β-Lapachone for 2 h every other day for 3 sequential treatments, followed by collection for genomic DNA extraction. (B)-(C) Analysis and categorization of hits from the sensitization screen. (B) Volcano plots summarizing gene significances based on sgRNA abundance changes between β-Lapachone treatment versus DMSO treatment. Left, A549 cells were treated with either DMSO or 2.0 μM β-Lapachone. Right, HCC15 cells were treated with either DMSO or 4.0 μM β-Lapachone. Treatment strategy was as described in (A). p-values were calculated by MAGeCK. Selected screen hits are highlighted and categorized by color. (C) Selected screen hits organized by category and color coded to correspond to (B). (D) Validation of SOD2. A549 cells and HCC15 cells were infected with virus encoding for either a non-targeting control sgRNA or sgRNAs against SOD2. Left, cells were exposed to either DMSO or escalating concentrations of β-Lapachone for 2 h, after which medium was replaced and remaining cell quantity was assessed 48 h after treatment using crystal violet staining. Right, western blot analyses of SOD2 and HSP90 (loading control).

In this study we focus on the results of sensitization screen. We found that enzymes in the pentose phosphate pathway (G6PD, PGLS, PGD) were highly represented among the enzymes whose loss sensitize cells to β-Lapachone, which is consistent with their reported role in NADPH generation for ROS detoxification [24]. Of note, we also find sgRNAs targeting thioredoxin reductase 1 (TXNRD1) and thioredoxin (TXN) among the top hits, consistent with our previous findings that the thioredoxin system plays a role in resistance of KEAP1/NRF2 mutant cells to β-Lapachone [25]. Interestingly, the mitochondrial localized thioredoxin reductase 2 (TXNRD2) did not score as a hit. TXNRD1 is a selenoprotein whose translation requires the selenocysteine biosynthesis pathway and we also found that selenophosphate synthetase 2 (SEPHS2) scored in both sensitivity screens, highlighting the role of selenoproteins in ROS detoxification in NSCLC. Within the thioredoxin-dependent antioxidant system, both the cytosolic (PRDX1) and mitochondrial (PRDX3) peroxidases scored. Further, in agreement with our previous report demonstrating that inhibition of glutathione synthesis does not sensitize NSCLC cells to β-Lapachone [25], there was a lack of glutathione-dependent enzymes in the hit list, and only small fold-changes for the glutathione synthesis enzymes GCLM and GCLC (Supplementary Table 1). Interestingly, we found glutathione reductase (GSR) scored strongly (Figs. 1B and 1C), suggesting that oxidized glutathione may have a specific role in antagonizing cell viability following β-Lapachone treatment. To validate these findings, we infected cells with lentivirus encoding for new hit-targeting sgRNAs or a non-targeting control sgRNA and confirmed protein loss by western blot. We found that loss of TXNRD1 or GSR strongly sensitized cells to β-Lapachone treatment (Fig. S1D), while TXNRD2 loss did not (Fig. S1E), consistent with the screen results.

While many of our hits were components of the glutathione and thioredoxin antioxidant systems, or the pentose phosphate pathway, the mitochondrial superoxide dismutase SOD2 scored as the sole superoxide metabolizing enzyme that was a top hit in both of the sensitization screens (Figs. 1B, S1C). We previously reported that inhibition of the cytosolic isoform superoxide dismutase SOD1 increases sensitivity to β-Lapachone [25], but the role of SOD2 was not explored. We further validated the effects of SOD2 inhibition on β-Lapachone efficacy. We infected both HCC15 and A549 cells with lentivirus encoding for sgRNA against SOD2 or a non-targeting control sgRNA. Consistent with the screen results, depletion of SOD2 markedly increased the sensitivity of A549 and HCC15 cells to β-Lapachone (Fig. 1D).

### 2.2 SOD2 inhibition increases β-Lapachone sensitivity in NSCLC

SOD2 is exclusively localized in the mitochondria matrix, which is a site of superoxide (O_2_•^−^) production by the electron transport chain (ETC). As a result, SOD2 is the principal mitochondrial O_2_•^−^ scavenger and an indispensable enzyme to maintain the redox balance in mitochondria. To understand the role of SOD2 in β-Lapachone sensitivity, we first examined whether NRF2/KEAP1 mutant cells have elevated SOD2. While several studies have suggested SOD2 is an ARE-responsive antioxidant gene, there is no direct evidence of NRF2 binding to the SOD2 ARE. Therefore, we examined the protein levels of SOD2 across our panel of NSCLC cells to evaluate whether KEAP1 wild-type (KEAP1^WT^) cells have higher SOD2 expression levels upon NRF2 activation. We treated KEAP1^WT^ cell lines with the KEAP1/NRF2 interaction inhibitor Ki-696 to stabilize NRF2. As expected, NRF2 activation induced the expression of well-characterized NRF2 targets, including GSR (Fig. 2A) and TXNRD1 (Fig. S2A). However, we did not observe the induction of SOD2. We further interrogated whether NRF2 deletion decreased SOD2 protein levels by infecting KEAP1 mutant (KEAP1^MUT^) cells with sgRNA against NRF2 or a non-targeting control sgRNA. Depletion of NRF2 reduced protein levels of well-characterized NRF2 targets, including GSR, NQO1 and TXN (Figs. 2B, S2B), but did not decrease SOD2. Collectively, these results demonstrate that SOD2 is not regulated by NRF2 in NSCLC cells. In contrast, SOD2 was previously shown to be induced γ-irradiation [30, 31], suggesting DNA damage or free radicals may induce its expression. Importantly, we observed that SOD2 protein levels were upregulated following β-Lapachone treatment in both KEAP1^WT^ and KEAP1^MUT^ cell lines (Fig. 2C). These results suggest that SOD2 induction is a protective response to β-Lapachone.

**Fig. 2.**
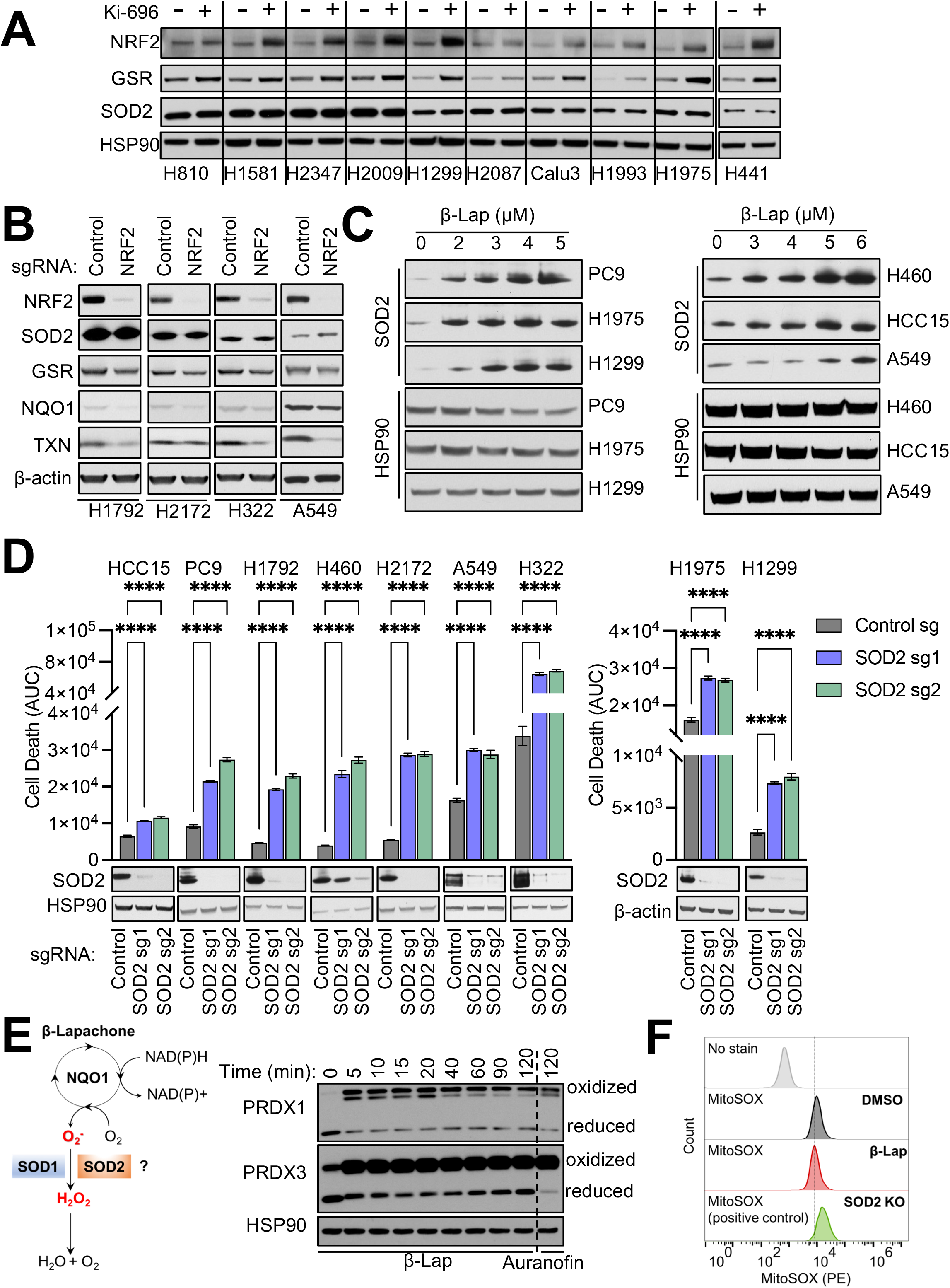
β-Lapachone increases SOD2 expression and dependence in NSCLC cells. (A-B) NRF2 does not regulate SOD2 expression. (A) Immunoblot of NRF2, GSR, and SOD2 expression in a panel of KEAP1/NRF2^WT^ cells pre-treated with 100 nM KI-696 or vehicle (DMSO) for 48 h. HSP90 was the loading control. (B) Immunoblot of NRF2, GSR, SOD2, NQO1 and TXN expression in a panel of KEAP1^MUT^ cells with Cas9 expression infected with virus encoding for either a non-targeting control sgRNA or sgRNA against NRF2. β-actin is used as the loading control. (C) β-Lapachone induces SOD2 expression. Immunoblot of SOD2 in a panel of KEAP1^WT^ cells (Left) and a panel of KEAP1^MUT^ cells (Right). Cells were treated with either DMSO or escalating concentrations of β-Lapachone for 2h, after which medium was replaced and protein was extracted 24 h after treatment. HSP90 was used as the loading control. (D) SOD2 deletion increases β-Lapachone cytotoxicity. Top, control sgRNA or SOD2 sgRNA expressing cells were treated with DMSO or β-Lapachone for 2 h, after which the medium was replaced. Left, NRF2 hyperactive NSCLC cells (β-Lapachone = 4 μM). NRF2 amplified: PC9; KEAP1 mutant: HCC15, H1792, H460, H2172, A549, H322. Right, KEAP1/NRF2^WT^ NSCLC cells: H1975, H1299 (β-Lapachone = 3 μM). Cell death was determined by Incucyte analysis of Sytox Green staining over 72 h, followed by normalization to cell density. Area under the curve (AUC) calculations are presented. Bottom, immunoblot analyses of SOD2 and HSP90 (loading control). Data are shown as mean ± SD. ****p < 0.0001. Two-way ANOVA with Dunnett’s multiple comparison test was used for statistical analyses. (E) Left, schematic representation of the potential role of SOD1 and SOD2 in the detoxification of β-Lapachone-induced ROS. Right, redox immunoblot analysis of the oxidation state of PRDX1 and PRDX3 in parental A549 cells following treatment with DMSO or 2 μM β-Lapachone for the indicated time, or 6 μM Auranofin for 120 min. HSP90 was the loading control. (F) β-Lapachone does not generate mitochondrial superoxide. A549 cells were treated with DMSO or 3 μM β-Lapachone for 10 min and mitochondrial superoxide assayed with MitoSOX Red. SOD2 KO A549 cells were used as a positive control.

We next directly evaluate the consequence of SOD2 deletion in response to β-Lapachone in NSCLC cells. We exposed a panel of NSCLC cell lines with NRF2 hyperactivation to vehicle control or β-Lapachone for 2 hours and monitored viability over 72 hours using the fluorescent dye Sytox Green, which stains the nuclei of dead cells. Cumulative cell death was calculated from the area under the curve (AUC), thereby facilitating comparisons between cell lines with different genetic modifications as we previously described [32]. Consistent with the results from the screen and crystal violet staining, SOD2 deletion with two independent sgRNAs significantly increased cell death in response to an escalating concentration of β-Lapachone treatment (Fig. S2C). We next expanded our examination to a panel of NSCLC cell lines with NRF2 hyperactivation. Upon challenging with β-Lapachone, cells with SOD2 deletion demonstrated significantly more cell death (Fig. 2D). However, these findings were not exclusive to NRF2 hyperactive cells as SOD2 deletion also significantly increased the death of KEAP1^WT^ cells following β-Lapachone exposure (Fig. 2D). Collectively, these data demonstrated that SOD2 deletion increased cell death in response to β-Lapachone treatment in NSCLC.

β-Lapachone is reduced by NQO1 to a labile hydroquinone which spontaneously re-oxidizes to β-Lapachone, thereby generating O_2_•^−^ and oxidative stress. Given the unique role of SODs in catalyzing the dismutation of O_2_•^−^ (Fig. 2E), we next examined whether the increased β-Lapachone sensitivity upon SOD2 deletion is due an inability to detoxify β-Lapachone-derived O_2_•^−^. To answer this question, we first determined whether β-Lapachone treatment induces mitochondrial oxidative stress. We assessed the oxidization state of both the cytosolic (PRDX1) and mitochondrial (PRDX3) peroxiredoxins by redox Western blotting. Because H_2_O_2_ detoxification by PRDX induces disulfide bond-mediated dimerization, which is subsequently reduced by TXN to restore PRDX antioxidant function, the ratio of PRDX monomers to dimers can be used as a surrogate marker for oxidative stress [33]. We found that β-Lapachone treatment resulted in a rapid and substantial oxidation of both PRDX1 and PRDX3 within 5 min, indicating that β-Lapachone induces ROS in both cytosolic and mitochondrial fractions (Fig. 2E). However, β-Lapachone-induced oxidation of PRDX3 could be a consequence of H_2_O_2_ diffused from the cytosol, where β-Lapachone metabolism by NQO1 generates O_2_•^−^, which is subsequently metabolized to H_2_O_2_. To test whether β-Lapachone can directly generate O_2_•^−^ in the mitochondria, we monitored mitochondrial superoxide levels using the MitoSOX Red probe. We found that while SOD2 deletion elevated mitochondrial O_2_•^−^, β-Lapachone treatment did not (Fig. 2F), nor did it elevate mitochondrial O_2_•^−^ in SOD2 deficient cells (Figs. S2D, S2E). These results indicate β-Lapachone does not generate O_2_•^−^ in mitochondria, but rather induces mitochondrial oxidative stress via the diffusion of H_2_O_2_.

### 2.3 SOD2 loss leads to a defect in mitochondrial ATP generation upon β-Lapachone treatment

Given that β-Lapachone does not generate O_2_•^−^ in mitochondria, it was surprising that SOD2 would play a protective role against β-Lapachone cytotoxicity. SOD2 plays a critical role in preventing mitochondrial oxidative damage, therefore we examined mitochondrial function following SOD2 loss. Using the Seahorse extracellular flux analyzer, we observed that the oxygen consumption rate (OCR) of SOD2-deficient A549 and HCC15 cells was significantly reduced compared to control cells (Fig. S3A). Notably, the spare respiratory capacity of SOD2-deficient cells was significantly lower (Fig. 3A), which is suggestive of a mitochondrial ATP generation defect upon an increase in energy demand. Indeed, when we examined cellular ATP levels, we found that SOD2-deficient cells failed to maintain ATP levels to that of the control cells following β-Lapachone treatment (Fig. 3B). However, total ATP levels do not reflect the contribution of ATP generation from individual compartments. Therefore, we performed a more specialized Seahorse-based protocol that permits the simultaneous analysis of mitochondrial ATP production and glycolytic ATP production. As expected, we found that SOD2-deficient cells exhibited significantly reduced mitochondrial ATP generation following β-Lapachone treatment, while glycolytic ATP was unchanged (Fig. 3B). These results indicate that SOD2 deletion results in impaired mitochondrial ATP production and suggest a bioenergetic crisis mediates loss of cell viability.

**Fig. 3.**
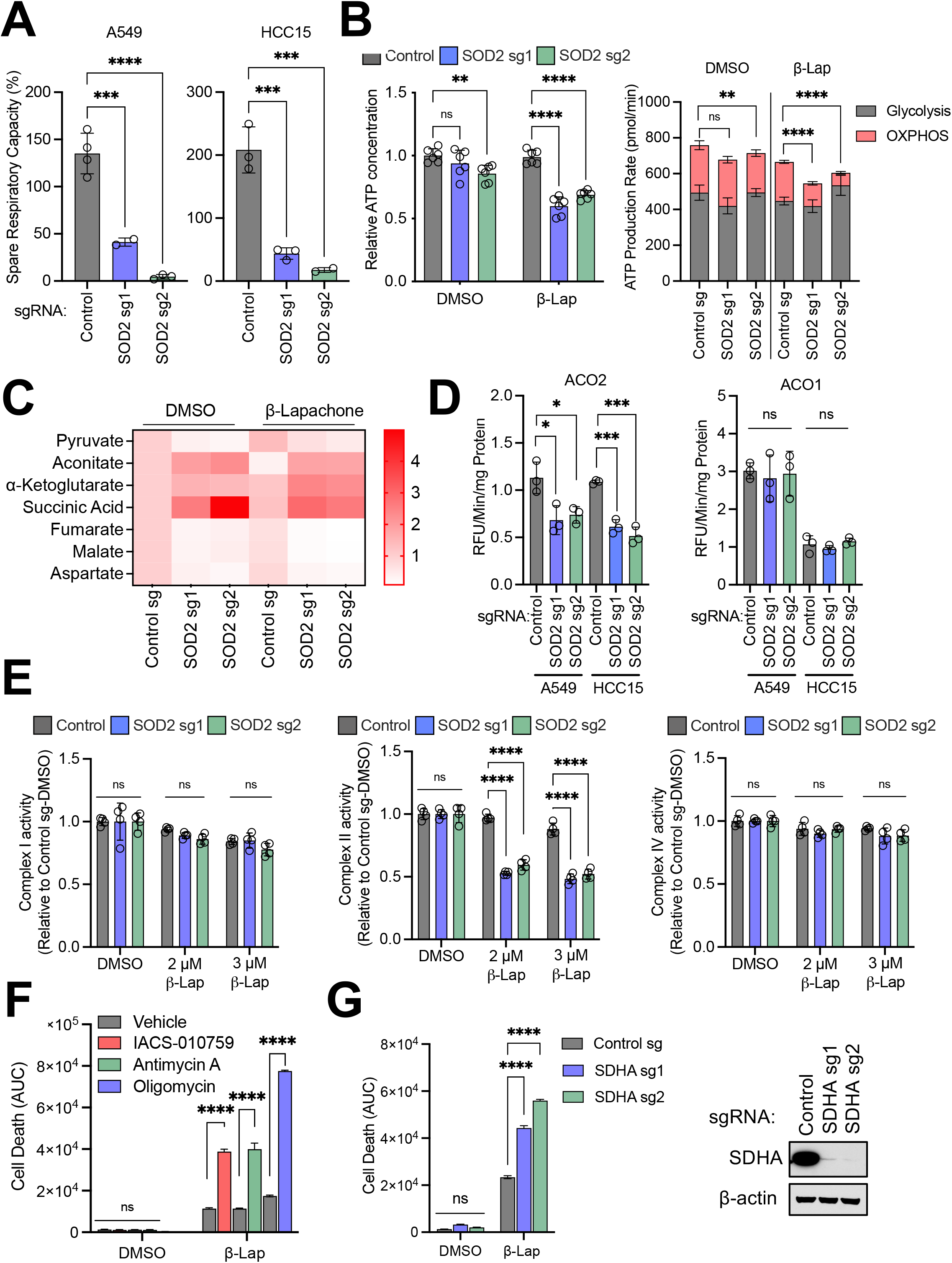
SOD2 loss leads to a defect in mitochondrial ATP generation upon β-Lapachone treatment. (A) Average spare respiratory capacity of A549 and HCC15 cells. Cells were previously infected with lentivirus encoding either control sgRNA or sgRNAs against SOD2. Student’s t test was used for statistical analyses. (B) Left, effect of SOD2 deletion on total cellular ATP levels. A549 control sgRNA and SOD2 sgRNA expressing cells were subjected to either DMSO or 2 μM β-Lapachone for 2 h. ATP levels were then measured using CellTiter-Glo assay and normalized to that of control cells treated with DMSO. Right, determination of the ATP production rate. Control and SOD2 sgRNA expressing cells were treated with DMSO or 2 μM β-Lapachone for 2 h, and the ATP production rate was measured using Agilent Seahorse XF Real Time ATP rate assay. Two-way ANOVA with Dunnett’s multiple comparison test was used for statistical analyses. *p < 0.05; **p < 0.01; ****p < 0.0001; ns, not significant. (C) Heatmap of relative abundances of TCA cycle metabolites in control and SOD2 sgRNA expressing A549 cells treated with DMSO or 2 μM β-Lapachone for 1.5 h. Data were normalized to the A549 control DMSO group (n = 3). (D) Mitochondrial (ACO2, Left) and cytosolic (ACO1, Right) aconitase activity of A549 and HCC15 cells following SOD2 deletion. One-way ANOVA with Dunnett’s multiple comparison test was used for statistical analyses. *p < 0.05; ***p < 0.01; ns, not significant. (E) Analysis of ETC complex I, II and IV activity following β-Lapachone treatment using Seahorse analysis with permeabilization. A549 cells subject to SOD2 deletion were treated with DMSO or the indicated concentrations of β-Lapachone for 2 h. (F) Mitochondrial ETC inhibition enhances β-Lapachone-induced cell death. A549 cells were treated with DMSO or 4 μM β-Lapachone for 2 h in combination with vehicle (DMSO) or the indicated mitochondrial ETC inhibitors, after which time the medium was replaced. Cell death was determined by Incucyte analysis of Sytox Green staining over 72 h, followed by normalization to cell density. Area under the curve (AUC) calculations are presented. Two-way ANOVA with Dunnett’s multiple comparison test was used for statistical analyses. Data are shown as mean ± SD. ****p < 0.0001. ns, not significant. (G) Left, control or SDHA sgRNA expressing A549 cells were treated with DMSO or 4 μM β-Lapachone for 2 h, after which the medium was replaced. Cell death was determined by Incucyte analysis of Sytox Green staining over 72 h, followed by normalization to cell density. Area under the curve (AUC) calculations are presented. Right, western blot analyses of SDHA and HSP90 (loading control). Data are shown as mean ± SD. Two-way ANOVA with Dunnett’s multiple comparison test was used for statistical analyses. ****p < 0.0001; ns, not significant.

To examine the mechanism underlying the decrease in mitochondrial function upon SOD2 loss, we performed liquid chromatography-high resolution mass spectrometry (LC-HRMS)-based metabolomics on control or SOD2 deleted A549 cells, in the presence of either vehicle control or β-Lapachone treatment. Analysis of TCA cycle metabolites revealed accumulation of substrates of Fe-S proteins, including succinate and aconitate, with loss of other TCA cycle intermediates including malate and fumarate (Fig. 3C). Increased oxidative stress can cause oxidation of mitochondrial electron transport chain (ETC) complexes containing iron-sulfur (Fe-S) clusters. NADH dehydrogenase (Complex I), succinate dehydrogenase (SDH; Complex II) and cytochrome reductase (Complex III) all contain Fe-S clusters, which are particularly sensitive to oxidation and subsequent inactivation by O_2_•^−^. To directly assess the impact of SOD2 on Fe-S protein function, we first assessed the activity of aconitase 2 (ACO_2_), a Fe-S protein of TCA cycle that is highly sensitive to oxidative inactivation. We found that SOD2 loss significantly reduced ACO_2_ activity in both A549 and HCC15 cells, while the cytosolic isoform aconitase 1 (ACO1) activity as unchanged (Fig. 3D). We next evaluated the activity of individual ETC complexes in permeabilized cells [33]. In the presence of Complex I substrates pyruvate and malate, the activity of Complex I was maintained to the level of control cells following SOD2 deletion in DMSO or β-Lapachone treated conditions. However, in the presence of succinate, SOD2 loss in combination with β-Lapachone exposure significantly reduced Complex II activity (Fig. 3E). As expected, SOD2 loss or β-Lapachone exposure did not alter the Complex IV activity, which does not contain Fe-S cluster, supporting that the loss of mitochondrial function following SOD2 loss is Fe-S cluster-dependent (Fig. 3E). Collectively, these data indicate that SOD2 deletion leads to a defect in mitochondrial Fe-S cluster containing protein function, thereby compromising the ETC and impairing mitochondrial ATP production.

To determine whether loss of mitochondrial function plays a causal role in sensitization to β-Lapachone, we co-treated cells with mitochondrial complex inhibitors and β-Lapachone to mimic the effect of SOD2 deletion. As expected, 2 hour co-treatment of β-Lapachone with inhibitors of ETC complex I or complex III both demonstrated a synergistic effect on cell death (Fig. 3F). Similarly, deletion of the ETC complex II component SDHA recapitulated the phenotype of SOD2 deletion and increased β-Lapachone efficacy (Fig. 3G). Notably, co-treatment of β-Lapachone with the complex V inhibitor oligomycin, which directly inhibits mitochondrial ATP synthesis, demonstrated the most drastic induction of cell death (Fig. 3F), further supporting the role of loss of ATP synthesis in β-Lapachone-induced cell death downstream of SOD2 loss. Together, these data indicate that SOD2 deletion leads to impaired mitochondrial ATP production, resulting in increased cell death following β-Lapachone treatment in NSCLC cells.

### 2.4 SOD2 loss lowers the NADPH/NADP+ ratio following β-Lapachone treatment

Our screen identified PPP components (G6PD, PGLS, and PGD) as the top hits strongly depleted from β-Lapachone treated cells, which is consistent with their annotated role as the primary source of NADPH generation. NADPH is an important redox cofactor that is served as the electron donor for GSR and TXNRD to reduce GSH and TXN, respectively. As the majority of our top hits converge on NADPH, either through its generation or utilization, we examined whether the effects of SOD2 deletion involved NADPH. Indeed, we observed a shift toward a lower NADPH/NADP+ ratio upon SOD2 depletion. Importantly, the reduction of NADPH/NADP+ ratio was greater in SOD2 deficient cells under β-Lapachone treatment, consistent with NADPH being consumed in oxidative stress conditions (Fig. 4A).

**Fig. 4.**
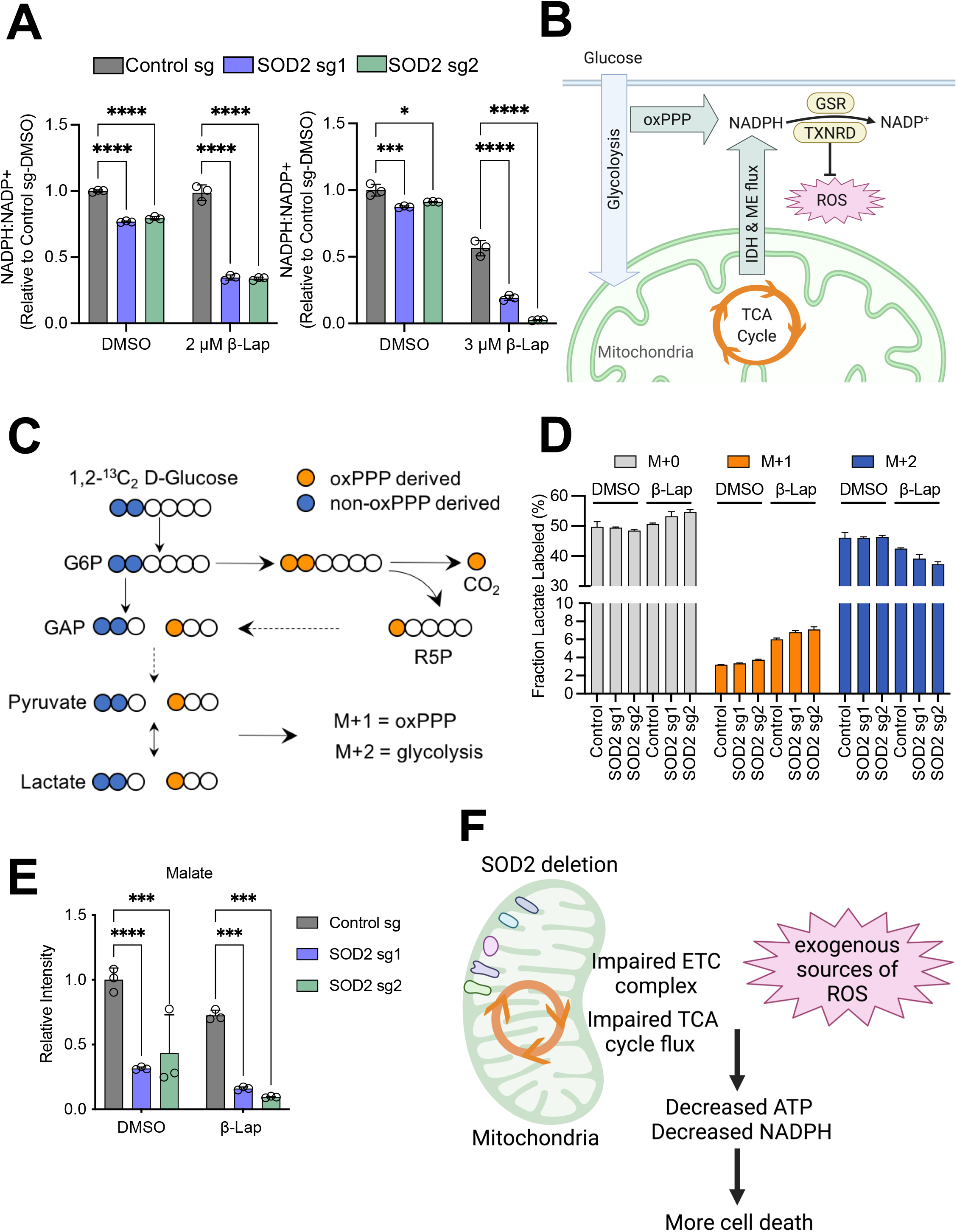
SOD2 loss lowers the NADPH/NADP+ ratio following β-Lapachone treatment. (A) NADPH:NADP+ ratio measurement. A549 cells expressing control or SOD2 sgRNAs were treated with DMSO or the indicated concentration of β-Lapachone for 2 h prior to collection. The NADPH:NADP+ ratios were normalized to control DMSO treated cells. (B) Schematic depiction of NADPH metabolism in NRF2 hyperactivation cells. (C) Tracer strategy for the analysis of pentose phosphate pathway flux using [1,2-^13^C] glucose. Dark blue circles depict glycolytic metabolites produced exclusively by glycolysis (M+2 labeling); orange circles depict glycolytic intermediates arising from glucose that first passed through the oxPPP (M+1). This schematic was adapted from Ghergurovich JM, et al. Local production of lactate, ribose phosphate, and amino acids within human triple-negative breast cancer. Med (N Y). 2021 Jun 11;2(6):736-754. (D) M+0 (unlabeled), M+1 (PPP-derived) and M+2 (glycolysis derived) labeling of lactate from [1,2-^13^C] glucose (mean + SD; n = 3). A549 cells expressing either control or SOD2 sgRNAs were treated with DMSO or 2 μM β-Lapachone for 1.5 h before metabolite extraction. See also Figure S4 (A-B). (E) Relative abundance of malate in A549 control and SOD2 sgRNA expressing cells treated with DMSO or 2 μM β-Lapachone for 1.5 h. Data was normalized to the control DMSO treated group. (n = 3). Two-way ANOVA with Dunnett’s multiple comparison test was used for statistical analyses. ***p < 0.001; ****p < 0.0001. (F) Model: SOD2 deletion upon excessive oxidative stress leads to reduction of ATP and NADPH, and ultimately accelerated cell death.

As glucose flux through G6PD and PPP is a substantial source of NADPH production (Fig. 4B), we questioned whether the reduced NADPH/NADP+ ratio was due to decreased glucose flux through the PPP in SOD2 deficient cells. To explore this possibility, we performed [1, 2 -^13^C] glucose tracing (Fig. 4C), and found that, upon β-Lapachone treatment, glucose carbon flux into PPP was elevated, as indicated by increased M+1 labeling of lactate (Fig. 4D), pyruvate (Fig. S4A) and Ribose-5-phostate (Fig. S4B). However, glucose carbon flux into the PPP was not influenced by SOD2 deletion (Figs. 4D, S4A, S4B), indicating the lower NADPH/NADP+ ratio upon SOD2 deletion is not a consequence of altered PPP flux. In addition to the PPP, Isocitrate dehydrogenase (IDH1 and IDH2) and malic enzymes (ME1 and ME3) represent other major sources of NADPH generation (Fig. 4B). Previous studies indicate that NRF2 activation induces cytoplasmic IDH1 and ME1, highlighting their role in maintaining the NADPH pool in NRF2 hyperactivated NSCLC cells [19, 23]. Cytoplasmic ME1 enzyme consumes the TCA intermediate malate in the generation of NADPH. We observed that in the absence of SOD2, malate was strikingly depleted, which became even more pronounced upon β-Lapachone treatment (Figs. 3C, 4E). We did not observe IDH1 or ME1 protein levels change upon SOD2 deletion or β-Lapachone treatment (Fig. S4C). Therefore, it is likely that the depletion of TCA intermediates upon SOD2 deletion, especially malate, leads to a deficit of NADPH in response to β-Lapachone, and consequently, impaired ROS detoxification. Altogether, these data support a model where defects in mitochondrial ATP maintenance and NADPH production resulting from SOD2 deletion contribute to the sensitization of NSCLC cells to β-Lapachone (Fig. 4F).

## DISCUSSION

In this study we conducted antioxidant enzyme gene focused CRISPR screens to comprehensively identify key antioxidant enzymes that mediates the resistance to β-Lapachone induced oxidative stress in both A549 (adenocarcinoma) and HCC15 (squamous cell carcinoma) cells. In agreement with our prior publication [25], our screens demonstrated the critical role played by the TXN-dependent antioxidant system, and gives further specificity to which TXN-dependent enzymes are required for H_2_O_2_ detoxification. Importantly, our screen results support the notion that the TXN-dependent antioxidant system plays a more dominant role than the GSH-dependent antioxidant system in combating β-Lapachone induced ROS. Although the expression of these antioxidants is commonly transcriptionally regulated by NRF2 [14], PRXs are in high abundance and therefore, account for the largest percentage of H_2_O_2_ detoxification in cells [34, 35]. These results align with our observation that none of the GPXs were among the hits in the sensitization screen.

While our screen hits demonstrated the important role of H_2_O_2_ scavenging in combating β-Lapachone cytotoxicity, we also uncovered a surprising role for mitochondrial SOD2 in protection against cell death in both NRF2/KEAP1^WT^ and NRF2/KEAP1^MUT^ NSCLC cells. Importantly, our results suggest that SOD2 inhibition may sensitize cells to other ROS-generating therapeutics beyond β-Lapachone and its analogs. In agreement with this, a paraquat (PQ) based negative-selection CRISPR-based screen identified SOD2 as one of the hits strongly depleted from surviving population of paraquat treated Jurkat cells [36]. More recently, a set of CRISPR screens in RPE1 cells treated with different genotoxic agents illustrated the cellular response to DNA damage, identifying SOD2 as one of the top hits that is strongly depleted upon potassium bromate (KBrO3) treatment [37]. Both PQ and KBrO_3_ generate intracellular ROS, but through mechanisms of action distinct from β-Lapachone. PQ exists as a dication (PQ^2+^), which can accept electron from reducing equivalents, and this process is suggested to occur in both the cytosol and mitochondria. In the presence of O_2_, reduced PQ^2+^ converts O_2_ into O_2_•^−^ [36]; where KBrO3 induces intracellular ROS via generation of hydroxyl radicals of bromine or its oxides [38, 39]. Collectively, these findings highlight SOD2 as a unique target mediating the therapeutic ROS detoxification.

Our study demonstrates that the sensitization of SOD2 deficient cells to oxidative stress is associated with a defect in mitochondrial ATP maintenance. SOD2 catalyzes the dismutase reaction of O_2_•^−^ to H_2_O_2_ and oxygen, which is essential for protecting mitochondria from oxidative damage [40]. Previous work demonstrated cells lacking SOD2 exhibited elevated state levels of O_2_•^−^ and reduced SDH activity, likely due to inactivation of Fe-S clusters [41–44]. Moreover, it has been shown that disruption of SOD2 gene in human embryonic kidney 293 and human erythroleukemia cell lines resulted in decreased SDH activity and succinate acumination [45]. These findings are in line with our observations in NSCLC cell lines, supporting our findings that SOD2 loss promotes a defect in mitochondrial ATP generation. Importantly, our results indicate that development of compounds targeting mitochondrial ATP production may be a promising approach to sensitize tumor cells to ROS-generating therapies.

Our work also indicates a broader role for SOD2 in oxidant defense through the support of NADPH production in NRF2 hyperactive cells. Prior studies suggest that the oxidative branch of the PPP is the largest contributor to the cellular NADPH pool, while ME1 and IDH1 serve as a backup, with ME1 superior to IDH1, when there is a NADPH demand [18]. NRF2 also regulates ME1 and IDH1 in addition the PPP [19, 23, 46–48], suggesting that NRF2 hyperactive cells have increased capacity to maintain NADPH pools with TCA cycle metabolites [21]. We find that SOD2 loss leads to depletion of TCA cycle metabolites levels under oxidative stress, including malate, as well as the depletion of NADPH that was not attributable to a change in PPP activity. These findings suggest that depletion of TCA cycle intermediates impairs alternative NADPH-generating pathways such as ME1 and IDH1. We also found several PPP enzymes as top hits in our screens, which was expected based on their role in NADPH generation [18]. NRF2 activation promotes PPP enzyme expression to maintain high antioxidant capacity and nucleotide synthesis, supporting redox defense and biosynthetic needs of cancer cells [19, 49]. Very recently, PPP enzymes were identified as CRISPR dropout hits in KEAP1^MUT^ A549 and H1439 3D spheroid growth [22]. Consistently, a CRISPR/Cas9 genetic screen of metabolic genes identified G6PD as the synthetic lethal gene with KEAP1 mutation [23]. While these two studies highlight that NRF2 hyperactive cancer cells depend on PPP enzymes for proliferation, our screen results highlight that NRF2 hyperactivation cancer cells depend on PPP enzymes for excessive ROS detoxification, but while the PPP is necessary for ROS detoxification, it may not be sufficient in the context of impaired mitochondrial function.

We note our work provides only a partial view of the oxidative stress response to β-Lapachone. Indeed, our antioxidant enzyme-focused CRISPR screen library does not cover the entirety of human antioxidant enzyme genes. For instance, the cytosolic dismutase SOD1, malic enzyme 2 (ME2), CoQ oxidoreductase FSP1, and methylenetetrahydrofolate dehydrogenases 1 and 2 (MTHFD1 and MTHFD2) are not included in our library. While this library allowed us to profile the majority of the key components of β-Lapachone cytotoxicity, a more comprehensive library including both antioxidant and metabolic enzymes would provide power for an entire view of their architecture in combating ROS. Another limitation of this study is that we used A549 and HCC15 cells as models to represent the NSCLC response to β-Lapachone. Although these two extensively used NSCLC cell lines were highly consistent in their results, there are likely cell line specific responses to β-Lapachone that reflect distinct biology and cell of origin. Finally, β-Lapachone induced acute toxicity in red blood cells [50–52] preclude us from further assessing SOD2 inhibition *in vivo*. It will be fascinating to assess SOD2 inhibition whenever a tolerable version of β-Lapachone analog becomes available.

Taken together, our results represent a rich resource for the study of oxidative stress and highlight SOD2 as a unique antioxidant enzyme that could be exploited for therapeutic intervention in NSCLC.

## ACKNOWLEDGEMENTS

This work is supported by a Florida Department of Health Bankhead-Coley research program (9BC07) to G.M.D. This work has also been supported by the Lung Cancer Center of Excellence and Genomics and Proteomics/Metabolomics Core Facilities at the Moffitt Cancer Center, an NCI designated Comprehensive Cancer Center (P30-CA076292). We thank Dr. Stephen J. Elledge and Dr. Richard Olalekan Adeyemi for providing us the library. We thank Dr. David Boothman and Dr. Naveen Singh for providing us β-Lapachone. We thank Dr. Florian Karreth and Dr. Ana Gomes for the helpful discussions.

## AUTHOR CONTRIBUTIONS

Conceptualization, C.J. and G.M.D.; Methodology, C.J., and G.M.D.; Investigation, C.J., N.P.W., N.P.F., Y.P.K., A.T., and M.T.; Writing – Original Draft, C.J. and G.M.D.; Writing – Review & Editing, N.P.W, Y.P.K, and M.T.; Funding Acquisition, G.M.D.; Supervision, C.J., G.M.D.

## MATERIALS AND METHODS

### Cell lines and reagents

Parental NSCLC cell lines were previously described (DeNicola et al., 2015). Cell lines were routinely tested and verified to be free of mycoplasma (MycoAlert Assay, Lonza). All lines were maintained in RPMI 1640 media (Hyclone or Gibco) supplemented with 5% FBS without antibiotics at 37 °C in a humidified atmosphere containing 5% CO_2_ and 95% air. Lenti-X 293T cells were obtained from Clontech, and maintained in DMEM media (Hyclone or Gibco) supplemented with 10% FBS. Cell lines with stable expression of *Streptococcus pyogenes* Cas9 gene were generated by lentiviral transduction, followed by blasticidin selection as reported [53]. For selection of transduced cells, puromycin was added at the concentration of 1 μg/mL. Blasticidin was used at the concentration of 4 μg/mL.

### CRISPR Screen

We utilized an oxidative stress gene focused sgRNA library provide by Dr. Steven Elledge to identify genes whose loss sensitize KEAP1 mutant NSCLC cells to a sublethal concentration of β-Lapachone; and to identify genes which are required for β-Lapachone-induced cell death. The oxidative stress gene focused library was constructed in a lentiCRISPR V2 vector with an additional F + E modification in the tracer as described [54], which targets 139 genes and contains a pool of 1668 total sgRNA. For the CRISPR-based screens, 6 million Cas9+ A549 or HCC15 cells were spinoculated in the presence of 4 μg/μL polybrene at 300 g for 2 hours with the oxidative stress gene focused library at a multiplicity of infection (MOI) of 0.3 to limit co-transduction. Tissue culture plates were then returned to the incubator for additional 6 hours, followed by replenishment with fresh cell culture medium. Transduced cells were selected with 1 μg/mL of puromycin at 48 hours post transduction. Cells were then passaged every 72 hours, maintaining at least 5.6 million cells per library after each passage to keep adequate complexity. At day 5 post-puromycin selection, the positive- and negative-selection β-Lapachone screens were initiated. For the positive-selection screen, Cas9+ A549 cells were treated with DMSO or 2.5 μM β-Lapachone every other day for a total of 3X treatment; and Cas9+ HCC15 cells were treated with DMSO or 5.5 μM β-Lapachone every other day for a total of 3X treatment. For the negative-selection screen, Cas9+ A549 cells were treated with DMSO or 2.0 μM β-Lapachone every other day for a total of 3X treatment; and Cas9+ HCC15 cells were treated with DMSO or 4.0 μM β-Lapachone every other day for a total of 3X treatment. Per treatment, the media was replaced with fresh media containing the indicated concentrations of β-Lapachone or vehicle DMSO for 2 h, after which media was replaced by fresh medium. Once the screens were complete, the cells were harvested for genomic DNA extraction. The sgRNA inserts were PCR amplified, purified and sequenced on a MiSeq V3 (Illumina). The abundance of each sgRNA was obtained. The median log_2_ fold change in the abundance of all sgRNAs targeting a particular gene between DMSO and β-Lapachone treatment was then analyzed.

### Individual sgRNA CRISPR knockout analysis

For hit validation, individual sgRNA directed CRISPR/Cas9 knockout was performed as previously described [53]. In brief, sgRNAs of target genes were cloned into the pLentiGuide-puro (Addgene plasmid #52963) lentivector. Cas9+ NSCLC cells were then transduced by lentivirus encoding sgRNA targeting genes of interest. Transduced cells were then selected in the presence of puromycin. As a control, a non-targeting sgRNA was also cloned and transduced. On-target CRISPR effects were validated by immunoblot. Table S2 contains primers used for sgRNA expression vector construction.

### Western blot analysis

Lysates were prepared in RIPA lysis buffer (20 mM Tris-HCl [pH = 7.5], 150 mM NaCl, 1 mM EDTA, 1 % NP-40, 1% sodium deoxycholate) containing protease and phosphatase inhibitors. In brief, cell lysates were mixed with 6X sample buffer containing β-ME and were separated by SDS-PAGE electrophoresis using NuPAGE 4-12% Bis-Tris gels (Invitrogen), and transferred onto the 0.45 μm nitrocellulose membranes, blocked with 5% milk in TBST buffer and then probed with relevant primary antibodies at 4 °C overnight, followed by secondary antibody (Cell Signaling) incubation for 1 h at room temperature. Blots were then developed by incubation with ECL chemiluminescence or Clarity Western ECL Substrate (Bio-Rad) and film-based images were captured.

### Redox western blotting

Redox western blotting was performed as previously described [25]. In brief, cells were seeded in 6-well plates at 5 × 10^5^ cells/well. The next day, cell culture medium was replaced with medium containing DMSO, 3 μM β-Lapachone or 6 μM of auranofin (positive control) for the indicated amount of time. After treatment, the media was aspirated and cells were gently washed with 1 mL of ice-cold PBS. 1 mg/mL of bovine catalase was added to the alkylation buffer (40mM HEPES, 50mM NaCl, 1mM EGTA, complete protease inhibitors, pH 7.4) 30 min prior collection. Immediately prior sample collection, 200 mM of N-Ethylmaleimide (NEM) were added to the alkylation buffer and the lysis buffer was warmed to 42 °C for 1–2 min to dissolve the NEM. To lyse the cells, 200 μL of alkylation buffer were dispensed in each well, followed by 10 min incubation at room temperature. A solution of 10% CHAPS was added to the lysates to a final concentration of 1% CHAPS (20 μL/sample). Cell lysates were transferred to a 1.5 mL micro-centrifuge tube, samples were vortex and incubated on ice for further 30 min, followed by 15 min centrifugation at 13,000 x rpm, 4°C. The supernatant was transferred to a clean 1.5 ml microcentrifuge tube. These redox western samples were mixed with a 4X non-reducing buffer prior to separation by SDS-PAGE.

### Antibodies

The following antibodies were used: NRF2 (Cell Signaling Technologies, D1Z9C, Cat #12721), NQO1 (Sigma Aldrich, Cat # HPA007308), β-actin (Thermo Fisher, clone AC-15, Cat # A5441), α-tubulin (Santa Cruz, TU-02, Cat #sc-8035), SOD2 (Cell Signaling Technologies, Cat #13194S), Prdx3 (Abcam, Cat #ab73349), Prdx1 (Cell Signaling Technologies, D5G12, Cat# 50-191-580), HSP90 (Cell Signaling Technologies, Cat #4874S), ME1 (Thermo Fisher Scientific, Cat #PA5-21550), IDH1(Cell Signaling Technologies, Cat #8137S), SDHA (Cell Signaling Technologies, D6J9M, Cat #11998), GSR (Santa Cruz Biotechnology, Cat #sc-133245), TRXR1 (Cell Signaling Technologies,15140S), TRXR2 (Cell Signaling Technologies, Cat #12029S).

### Cell viability assays

NSCLC cells were seeded in 96-well plates at a density of 5,000 cells/well in a 100 μL final volume. The next day, the media was replaced with 200 μL of fresh media containing the indicated concentrations of β-Lapachone or vehicle (≤ 0.1% DMSO) for 2 h, after which the medium was replaced to 200 μL fresh medium. Cell viability was assessed 72 h after treatment with crystal violet staining. To stain surviving cells with crystal violet, cells were washed with ice-cold PBS, fixed with 4% paraformaldehyde, stained with crystal violet solution (0.1% Crystal Violet, 20% methanol), washed with H2O and dried overnight. Crystal violet was solubilized in 10% acetic acid for 30 min and the OD600 was measured. Relative cell number was normalized to vehicle DMSO treated cells.

### Dead Cell Measurement with IncuCyte

Cells were plated in black walled 96 well pates at a density of 2500 or 3,000 cells/well in 100 μL final volume. The next day, the medium was changed to 200 μL of experimental medium containing DMSO or β-Lapachone for 2h. In experiments additionally adding complex inhibitors, the experimental medium containing DMSO or β-Lapachone was further supplemented with 1 μM oligomycin, 1 μM antimycin A, or 1 μM IACS-010759 as indicated [33, 55]. After 2 h treatment, the medium was then removed and replaced with fresh medium containing 25 nM of Sytox Green. The number of dead cells and cell confluence were measured by the IncuCyte S3 live-cell analysis system (Essen BioScience, Ann Arbor, MI, USA) in a humidified tissue culture incubator at 37°C with 5% CO2. Data were acquired with a 10X objective lens in phase contrast and green fluorescence (Ex/Em: 460/524 nm, acquisition time: 400 ms) channels. Images were acquired from each well at 4-6 h intervals. Image and data processing were performed with IncuCyte S3 2018B, 2020A or 2021A software (Essen BioScience, Ann Arbor, MI, USA). Dead cell number was normalized to cell confluence [Number of Sytox Green positive cells/mm^2^/cell confluence (% of total image)]. The area under the curve (AUC) was calculated as the sum of the dead cell numbers at each time point.

### Sample preparation for non-targeted metabolite profiling

A549 cells expressing either control sgRNA or SOD2 sgRNAs were seeded in 6 well plates at a density of 5 ×10^5^ cell/well in a 2 mL final volume. On the following day, the media was replaced with fresh media containing the indicated concentration of β-Lapachone or DMSO for 1.5 h. Cells were quickly washed in cold PBS, and extracted in 0.5 mL 80% methanol (−80 ̊C, 15 min). The extracts were cleared by centrifugation, and the metabolites in the supernatant were directly analyzed by liquid chromatography-high resolution mass spectrometry (LC-HRMS).

### Stable isotope tracing

A549 cell infected with lentivirus encoding for sgRNAs against SOD2 or a non-targeting control cells were plated in 6 well dishes and pre-conditioned in RPMI medium containing dialyzed FBS (dFBS, 5%) overnight. The following day, the cells were quickly washed with 1 mL of glucose-free media, followed by feeding with 1, 2-^13^C_2_-Gluose containing medium (glucose-free RPMI + 5% dFBS + 2 g/L 1, 2-^13^C_2_-Gluose) supplemented with either vehicle control DMSO or 2 μM β-Lapachone. After 1.5 h, the medium was aspirated and the cells were quickly washed with ice cold PBS once. After aspirating the PBS, the cellular metabolites were extracted with 0.5 mL cold extraction solvent (80% MeOH: 20% H_2_O) at −80 °C for 15 min. After scraping, the metabolite extract was transferred into an Eppendorf tube and cleared by centrifugation (17,000 g, 20 min, 4 °C), followed by LC-MS analysis in negative mode.

### LC-MS Analysis

The LC-MS conditions were identical to previously established methods [56]. For the chromatographic metabolite separation, the Vanquish UPLC systems were coupled to a Q Exactive HF (QE-HF) mass spectrometer equipped with HESI (Thermo Fisher Scientific, Waltham, MA). The column was a SeQuant ZIC-pHILIC LC column, 5 μm, 150 x 4.6 mm (MilliporeSigma, Burlington, MA) with a SeQuant ZIC-pHILIC guard column, 20 x 4.6 mm (MilliporeSigma, Burlington, MA). Mobile phase A was 10 mM (NH_4_)_2_CO_3_ and 0.05% NH_4_OH in H_2_O while mobile phase B was 100% ACN. The column chamber temperature was set to 30 °C. The mobile phase condition was set according to the following gradient: 0-13 min: 80% to 20% of mobile phase B, 13-15 min: 20% of mobile phase B. The ESI ionization mode was negative. The MS scan range (m/z) was set to 60-900. The mass resolution was 120,000 and the AGC target was 3 x 10^6^. The capillary voltage and capillary temperature were set to 3.5 KV and 320 °C, respectively. 5 μL of sample was loaded. For the non-targeted metabolomics approach, the LC-MS peaks were automatically extracted and aligned using the Automated Feature Detection function of EL-Maven. After the normalization with the median value of the intensities of LC-MS peaks, the statistical analysis was conducted. For the isotope tracing experiment, the natural abundance isotope correction was performed using EL-Maven.

### Flow cytometry analysis

Control sgRNA expressing or SOD2 sgRNA expressing A549 cells were plated in 24 well dishes at 70,000 cells/well. The following day, the medium was aspirated and MitoSOX Red was added to the cells at a final concentration of 5 μM for 30 min. Cells were treated with DMSO or β-Lapachone for indicated amount of time in the presence of MitoSOX Red. After that, cells were washed with PBS, detached by trypsin and transferred with FACS buffer (PBS containing 0.5% BSA, 1 mM EDTA and 25 mM HEPES). Following brief centrifugation, the cell pellets were washed with FACS buffer and resuspend in 400 μL FACS buffer. The samples were analyzed on Accuri C6 flow cytometer (BD biosciences) using PE filter. The data was further analyzed with FlowJo (Ver 10.7.1) software (FlowJo).

### NADPH/NADP+ assay

A549 cells expressing either control sgRNA or SOD2 sgRNAs were seeded in 60 mm dishes at a 70% confluence in a 4 mL final volume. On the following day, the media was replaced with fresh media containing the indicated concentration of β-Lapachone or DMSO for 2 h. Cells were extracted and the NADPH/NADP+ ratio was measured using the NADP/NADPH-Glo Assay kit (Promega) according to the manufacturer’s instructions.

### Quantification of intracellular ATP

Cells were plated in 96-well plates at a density of 10,000 cells/well in 100 μL final volume. The next day, the media was replaced with fresh media containing the indicated concentration of β-Lapachone or DMSO for 2 h. An ATP standard curve was generated on the same plate on which the samples were assayed. CellTiter-Glo (Promega) was used to measure the ATP level in each well.

### Seahorse analysis of mitochondrial function

Measures of oxygen consumption was determined with a Seahorse XFe96 Analyzer (Agilent). General mitochondrial function was assessed according to the Seahorse XF Cell Mito Stress Kit protocol (Agilent). Briefly, cells were seeded at 40,000 per well in quadruplicate on an XFe96 microplate and allowed to settle down overnight. Immediately before the assay, cells were supplemented with 10 mM glucose and 1 mM glutamine and then sequentially challenged with 1 μM oligomycin, 1 μM of FCCP, and 1 μM each of antimycin A and rotenone. We assessed the individual ETC complex activity according to an established protocol [57]. Briefly, 40,000 cells were plated in quadruplicate on an XFe96 microplate and allowed to seed overnight. Immediately before assay, cells were overlaid with 175 μL of mitochondrial assay solution [33] supplemented with the Seahorse Plasma Membrane Permeabilizer (Agilent), 4 mM ADP (Sigma-Aldrich), and 10 mM sodium pyruvate (Sigma-Aldrich) with 1 mM malate (Sigma-Aldrich). Cells were then sequentially subjected to 2 μM rotenone (Sigma-Aldrich), 10 mM succinate (Sigma-Aldrich), 2 μM antimycin A (Sigma-Aldrich), and 10 mM ascorbate (Sigma-Aldrich) with 100 μM N,N,N’,N’-tetramethyl-ρ-phenylene diamine (Sigma-Aldrich). Glycolytic and mitochondrial ATP production rates were determined according to the XF Real-Time ATP Rate Assay Kit protocol (Agilent). Briefly, 40,000 cells were seeded overnight on an XFe96 microplate in prepared media mimicking Seahorse XF RPMI, pH 7.4 supplemented with 10 mM glucose (Sigma-Aldrich) and 2 mM glutamine (VWR). Cells were then overlaid with fresh media and sequentially subjected to 1 μM oligomycin and then concomitant 1 μM rotenone and antimycin A. Glycolytic and mitochondrial ATP production rates were calculated according to the “Quantifying Cellular ATP Production Rate Using Agilent Seahorse XF Technology” White Paper (Agilent).

### Aconitase Assay

Aconitase activity was determined according to a previously established protocol [33]. A549 or HCC15 cells subjected to SOD2 deletion or control cells were seeded the day before at 1*10^6 cells per dish in 10 cm dishes in triplicates. Cells were collected and resuspended in 250 μL of 50 mM Tris-HCl and 150 mM NaCl, pH 7.4. The cell suspension was homogenized with a dounce homogenizer and the homogenate spun down for 10 min at 10,000 g at 4°C. The supernatant was collected for ACO1 activity measurement. The pellet was then washed twice and resuspended in 100 μL of 1% Triton X-100 (Sigma-Aldrich) in 50 mM Tris-HCl, pH 7.4, to lyse the mitochondrial membrane. This fraction was then spun down for 15 min at 17,000 g at 4°C. The protein concentration for both cytosolic fraction and mitochondrial fraction was then determined by DC Protein Assay (Bio-Rad), and 175 μL of 100–500 μg/ml protein solution was generated with assay buffer (50 mM Tris-HCl, pH 7.4). 50 μL of this solution was transferred to triplicate wells of a black-walled 96-well fluorescence microplate already containing 55 μL of assay buffer. Next, 50 μL of a 4 mM NADP+ (Sigma-Aldrich), 20 U/ml IDH1 (Sigma-Aldrich) solution was added to each well. Finally, 50 μL of 10 mM sodium citrate (Sigma-Aldrich) was added to each well to initiate the assay. The plate was transferred to a fluorescence-compatible plate reader (Promega) to measure NADPH autofluorescence every minute over a period of an hour. This change in fluorescence over time is indicative of aconitase activity, where ACO converts the supplied citrate to isocitrate, which the supplied IDH1 then metabolizes in a reaction that generates NADPH.

### Statistical analysis

The statistical significance of CRISPR screen hits was calculated using the MAGeCK [58] by comparing sgRNA abundances in the DMSO treated cell population with levels in the β-Lapachone treated populations. In brief, MAGeCK evaluates the consistency across the sgRNAs targeting to the same genes and the reproducibility among triplicate experiments, to generate overall statistical significance prioritizing top gene hits. Multiple hypothesis testing was adjusted using the Benjamini-Hochberg method with false discovery rate (FDR) < 0.05 to filter significant screen hits.

Unless otherwise indicated, all bar graphs and line graphs represent the arithmetic mean representative of three independent experiments (n = 3), with error bars denoting standard deviations. Data were analyzed using two-tailed paired Student t test or analysis of variance (ANOVA) with the appropriate post-tests (Dunnett’s multiple comparison tests) using GraphPad Prism9 software. P values correlate with symbols as follows, ns = not significant, p > 0.05; * p < 0.05; ** p < 0.01; *** p < 0.001, **** p < 0.0001.

