## Supplemental Figures and Tables for "A CRISPR screen identifies redox vulnerabilities for KEAP1/NRF2 mutant non-small cell lung cancer"

### Figure S1

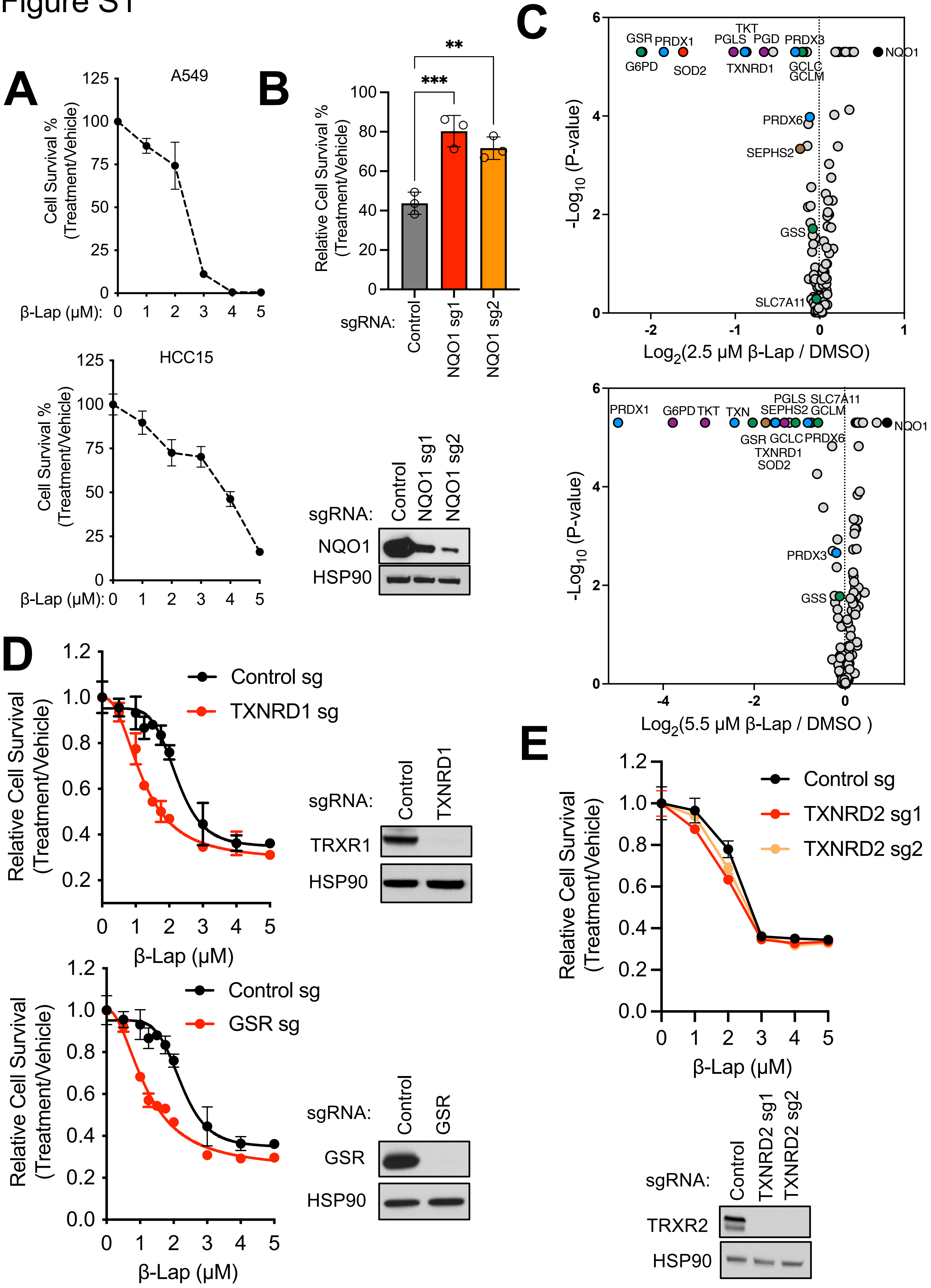

A
A549
Cell Survival % (Treatment/Vehicle)
β-Lap (μM):
HCC15
B
Relative Cell Survival % (Treatment/Vehicle)
sgRNA: Control NQO1 sg1 NQO1 sg2
NQO1
HSP90
C
-Log10 (P-value)
Log2(2.5 μM β-Lap / DMSO)
Log2(5.5 μM β-Lap / DMSO )
GSR PRDX1 TKT PGLS PGD PRDX3 NQO1 G6PD SOD2 TXNRD1 GCLC GCLM PRDX6 SEPHS2 GSS SLC7A11
PRDX1 G6PD TKT TXN PGLS SEPHS2 SLC7A11 GCLM GSR GCLC PRDX6 TXNRD1 SOD2 NQO1 PRDX3 GSS
D
Relative Cell Survival (Treatment/Vehicle)
Control sg
TXNRD1 sg
GSR sg
β-Lap (μM)
sgRNA: Control TXNRD1
TRXR1
HSP90
sgRNA: Control GSR
GSR
HSP90
E
Control sg
TXNRD2 sg1
TXNRD2 sg2
sgRNA: Control TXNRD2 sg1 TXNRD2 sg2
TRXR2
HSP90

**Fig. S1. CRISPR/Cas9 screens identify redox vulnerabilities of KEAP1 mutant NSCLC cells, related to Fig. 1.**

(A) Survival assays. A549 (Top) and HCC15 (Bottom) cells were exposed to either DMSO or escalating concentrations of β-Lapachone for 2 h every other day for 3 sequential treatments. After each treatment, medium was replaced. The remaining cell quantity was assessed 48 h after treatment using crystal violet staining. (B) NQO1 deletion promotes β-Lapachone resistance. Top, survival assays of A549 cells infected with virus encoding for sgRNAs against NQO1 or with virus encoding for a non-targeting control sgRNA. Cells were treated with vehicle (DMSO) or with 2.5 μM β-Lapachone for 2 h every other day for 3 sequential treatments. After each treatment, medium was replaced. The remaining cell quantity was assessed 48 h after treatment using crystal violet staining. One-way ANOVA statistical test was performed, followed by the multiple comparison Dunnett's test. *** $p < 0.0001$; ** $p < 0.001$. Bottom, western blot analyses of NQO1 and HSP90 (loading control). (C) Analysis of hits from the resistance screen. Volcano plots summarizing gene significances based on sgRNA abundance changes between β-Lapachone treatment versus DMSO treatment. A549 cells (Top) were treated with either DMSO or 2.5 μM β-Lapachone and HCC15 cells (Bottom) were treated with either DMSO or 5.5 μM β-Lapachone. Treatment strategy was as described in Fig 1(A). p-values were calculated by MAGeCK. Selected screen hits are highlighted. (D) Validation of TXNRD1 and GSR. A549 cells were infected with virus encoding for either a non-targeting control sgRNA or sgRNAs against TXNRD1 or GSR. Left, survival assays, cells were exposed to either DMSO or escalating concentrations of β-Lapachone for 2 h, after which medium was replaced and the remaining cell quantity was assessed 48 h after treatment using crystal violet staining. Right, western blot analyses of TXRX1, GSR and HSP90 (loading control). (E) TXNRD2 does not influence β-Lapachone sensitivity. A549 cells were infected with virus encoding for either a non-targeting control sgRNA or sgRNAs against TXNRD2 and treated with DMSO or β-Lapachone as described in (D). Right, western blot analyses of TXRX2 and HSP90 (loading control).

### Figure S2

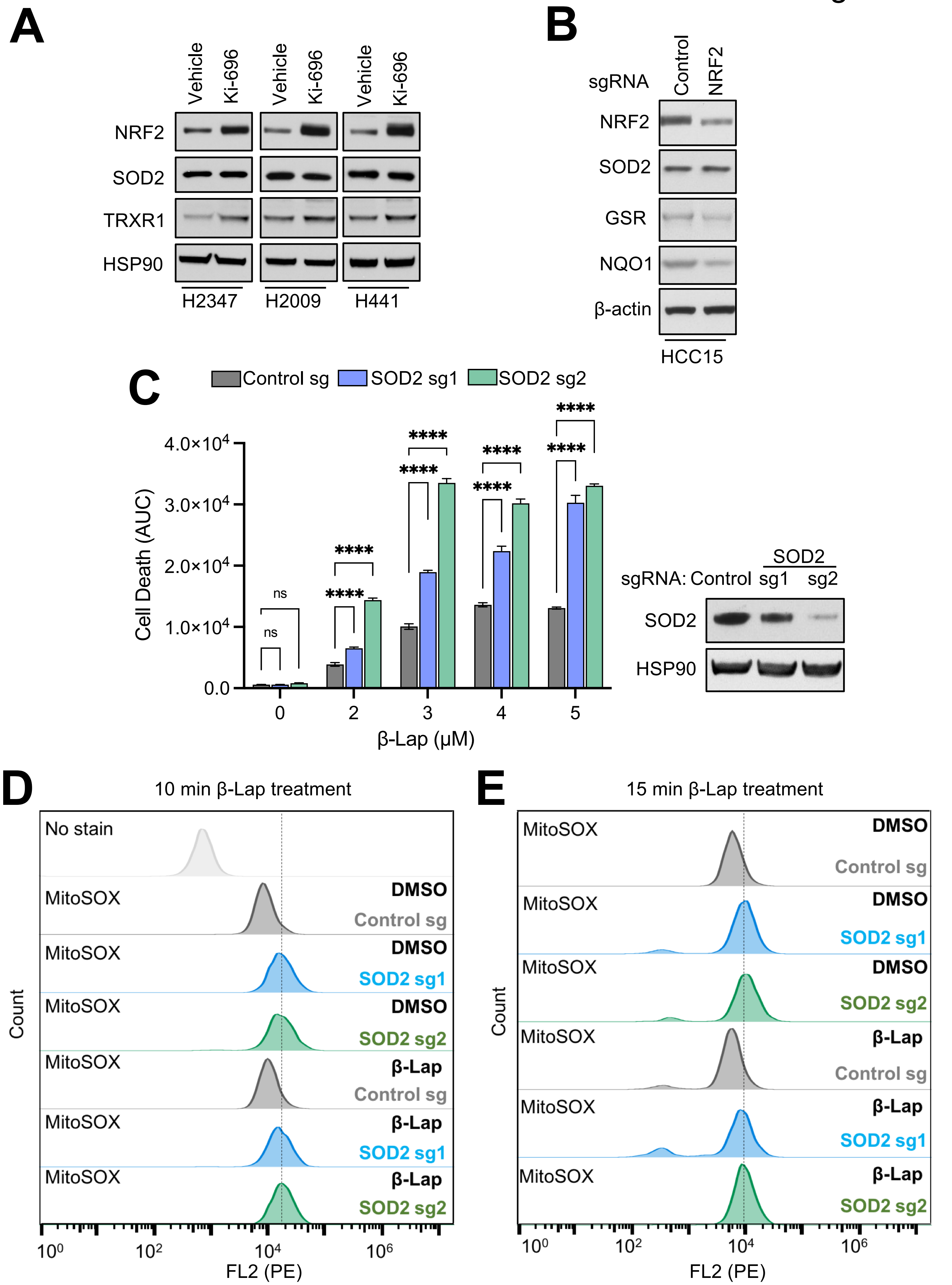

A
Vehicle
Ki-696
NRF2
SOD2
TRXR1
HSP90
H2347
H2009
H441
B
sgRNA
Control
NRF2
NRF2
SOD2
GSR
NQO1
β-actin
HCC15
C
Control sg
SOD2 sg1
SOD2 sg2
Cell Death (AUC)
4.0×10^4
3.0×10^4
2.0×10^4
1.0×10^4
0.0
ns
****
0
2
3
4
5
β-Lap (μM)
sgRNA: Control
SOD2
sg1
sg2
SOD2
HSP90
D
10 min β-Lap treatment
No stain
MitoSOX
DMSO
Control sg
SOD2 sg1
SOD2 sg2
β-Lap
Count
FL2 (PE)
E
15 min β-Lap treatment

**Fig. S2. β-Lapachone increases SOD2 expression and dependence in NSCLC cells, related to Fig. 2.**

(A-B) NRF2 does not regulate SOD2 expression. (A) Immunoblot of NRF2, TXRX1 and SOD2 expression in a panel of KEAP1/NRF2$^{WT}$ cells pre-treated with 100 nM KI-696 or vehicle (DMSO) for 48 h. HSP90 was used as the loading control. (B) Immunoblot of NRF2, GSR, and NQO1 expression in A549 cells with Cas9 expression infected with virus encoding for either a non-targeting control sgRNA or sgRNA against NRF2. β-actin was used as the loading control. (C) SOD2 deletion increases β-Lapachone cytotoxicity. Left, control sgRNA or SOD2 sgRNA expressing A549 cells were treated with DMSO or escalating concentrations of β-Lapachone for 2 h, after which the medium was replaced. Cell death was determined by Incucyte analysis of Sytox Green staining over 72 h, followed by normalization to cell density. Area under the curve (AUC) calculations are presented. Right, western blot analyses of SOD2 and HSP90 (loading control). Data are shown as mean ± SD. Two-way ANOVA with Dunnett's multiple comparison test was used for statistical analyses. ****$p < 0.0001$; ns, not significant. (D-E) Analysis of mitochondrial superoxide levels of A549 control sgRNA and SOD2 sgRNA expressing cells with MitoSOX Red. Cells were either treated with DMSO or 3 μM β-Lapachone for 10 min (D) or 15 min (E).

### Figure S3

**A**

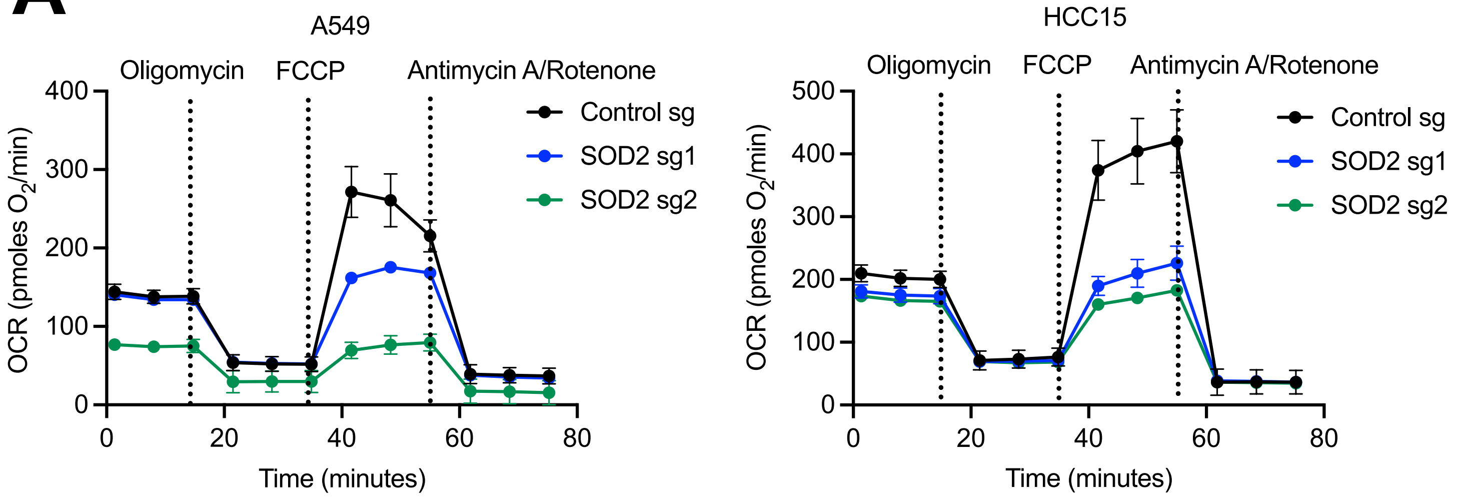

A549
Oligomycin
FCCP
Antimycin A/Rotenone
Control sg
SOD2 sg1
SOD2 sg2
OCR (pmoles O2/min)
Time (minutes)
HCC15
Oligomycin
FCCP
Antimycin A/Rotenone
Control sg
SOD2 sg1
SOD2 sg2
OCR (pmoles O2/min)
Time (minutes)

**Fig. S3. SOD2 loss leads to a defect in mitochondrial ATP generation upon β-Lapachone treatment, related to Fig 3.**

(A) Plots of OCR in A549 (left) and HCC15 (right) control sgRNA or SOD2 sgRNA expressing cells from Mito Stress test.

### Figure S4

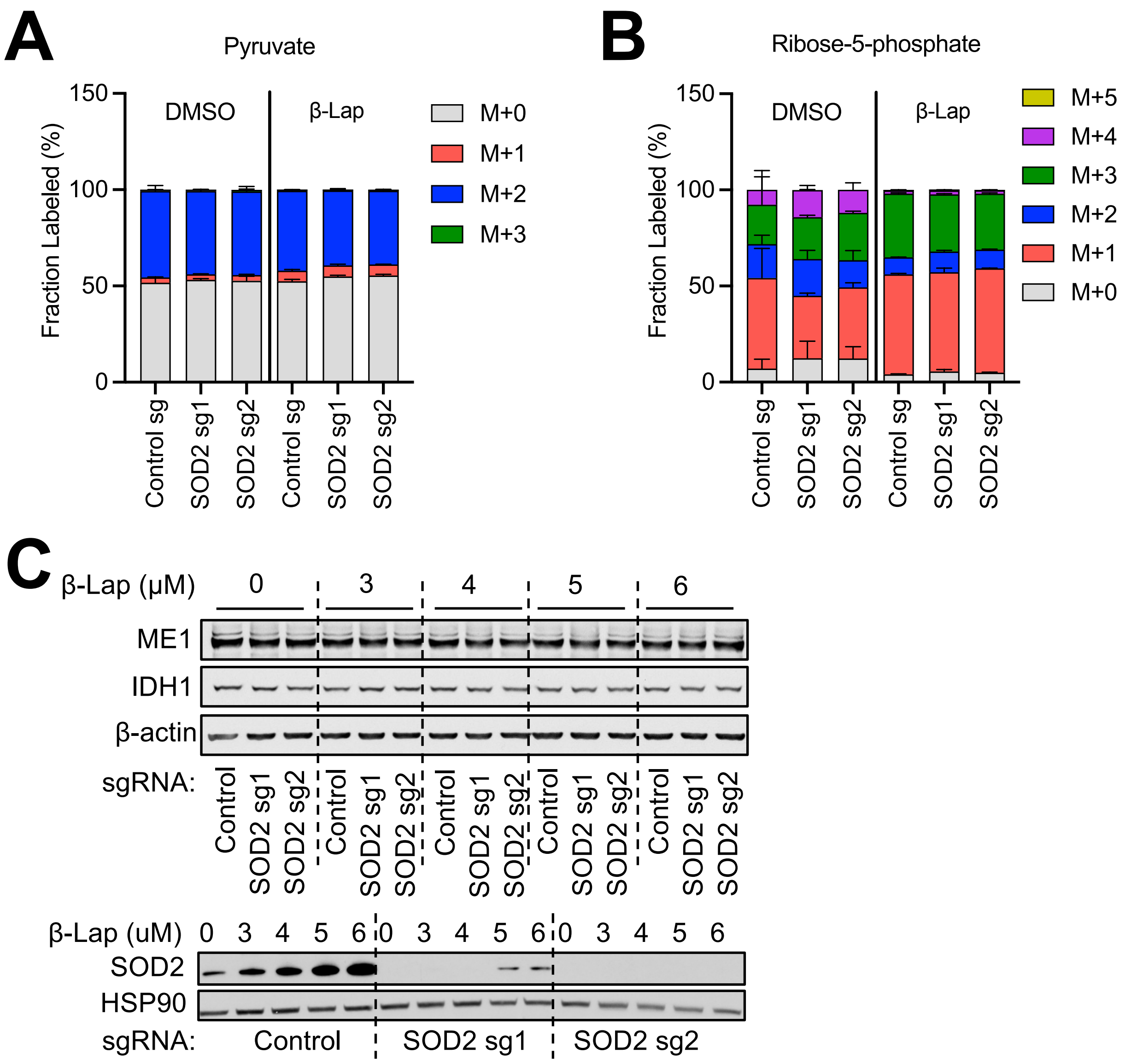
A
Pyruvate
Fraction Labeled (%)
150
100
50
0
DMSO
β-Lap
M+0
M+1
M+2
M+3
Control sg
SOD2 sg1
SOD2 sg2
Control sg
SOD2 sg1
SOD2 sg2
B
Ribose-5-phosphate
Fraction Labeled (%)
150
100
50
0
DMSO
β-Lap
M+5
M+4
M+3
M+2
M+1
M+0
Control sg
SOD2 sg1
SOD2 sg2
Control sg
SOD2 sg1
SOD2 sg2
C
β-Lap (μM)
0
3
4
5
6
ME1
IDH1
β-actin
sgRNA:
Control
SOD2 sg1
SOD2 sg2
β-Lap (uM)
0 3 4 5 6 0 3 4 5 6 0 3 4 5 6
SOD2
HSP90
sgRNA:
Control
SOD2 sg1
SOD2 sg2

**Fig. S4. SOD2 loss lowers the NADPH/NADP+ ratio following β-Lapachone treatment, related to Fig 4.**

(A-B) SOD2 loss does not alter pentose phosphate flux following β-Lapachone treatment. (A) M+0, M+1 and M+2 labeling of pyruvate from [1,2-$^{13}$C] glucose (mean + SD; n = 3); and (B) M+0, M+1, M+2, M+3, M+4, and M+5 labeling of Ribose -5-phosphate. A549 cells expressing either control or SOD2 sgRNAs were treated with DMSO or 2 μM β-Lapachone for 1.5 h before metabolite extraction. (C) Immunoblots of ME1, IDH1 and SOD2 in A549 cells expressing control or SOD2 sgRNAs treated with DMSO or escalating concentrations of β-Lapachone for 2 h. HSP90 and β-actin were used as the loading control.

**Table S1. β-Lapachone negative-selection CRISPR-based screen hits at FDR < 0.05 cutoff, related to Fig. 1.**

A549

| Gene Name | Negative Rank | FDR | $Log_2$ (Fold Change) [β-Lap=2.0 μM vs. DMSO] |
|---|---|---|---|
| G6PD | 1 | 5.80E-05 | -2.0243 |
| GSR | 2 | 5.80E-05 | -1.9038 |
| SOD2 | 3 | 5.80E-05 | -1.5075 |
| TKT | 4 | 5.80E-05 | -0.44258 |
| PRDX1 | 5 | 5.80E-05 | -1.6094 |
| PGLS | 6 | 5.80E-05 | -0.87924 |
| PGD | 7 | 5.80E-05 | -0.61794 |
| TXN | 8 | 5.80E-05 | -0.86327 |
| TXNRD1 | 9 | 5.80E-05 | -0.75126 |
| PRDX3 | 10 | 5.80E-05 | -0.23293 |
| DLD | 11 | 5.80E-05 | -0.3983 |
| GCLM | 12 | 5.80E-05 | -0.096505 |
| PFKP | 13 | 0.007351 | -0.11533 |
| PFKFB3 | 14 | 0.007351 | -0.095174 |
| AHCY | 15 | 0.007351 | -0.11722 |
| SEPHS2 | 16 | 0.009752 | -0.25243 |
| PRDX6 | 17 | 0.009913 | -0.1267 |
| GCLC | 18 | 0.025586 | -0.13502 |
| MGST1 | 19 | 0.025586 | -0.090667 |

HCC15

| Gene name | Negative Rank | FDR | $Log_2$ (Fold Change) [β-Lap=4.0 μM vs. DMSO] |
|---|---|---|---|
| G6PD | 1 | 3.50E-05 | -3.7514 |
| GSR | 2 | 3.50E-05 | -2.079 |
| TKT | 3 | 3.50E-05 | -2.684 |
| PRDX1 | 4 | 3.50E-05 | -5.0571 |
| HK1 | 5 | 3.50E-05 | -1.0777 |
| TXN | 6 | 3.50E-05 | -2.0633 |
| SOD2 | 7 | 3.50E-05 | -1.1124 |
| PGD | 8 | 3.50E-05 | -1.2926 |
| SEPHS2 | 9 | 3.50E-05 | -1.6546 |
| PGLS | 10 | 3.50E-05 | -1.0964 |
| GPI | 11 | 3.50E-05 | -0.89093 |
| GCLC | 12 | 3.50E-05 | -0.66228 |
| GCLM | 13 | 3.50E-05 | -0.32752 |
| PDHB | 14 | 3.50E-05 | -0.33631 |
| PRDX6 | 15 | 3.50E-05 | -0.27361 |
| PRDX3 | 16 | 3.50E-05 | -0.26539 |
| PDHA1 | 17 | 3.50E-05 | -0.32106 |
| TXNRD1 | 18 | 3.50E-05 | -1.2741 |
| ENO1 | 19 | 3.50E-05 | -0.26911 |
| SLC7A11 | 20 | 3.50E-05 | -0.21924 |
| GSS | 21 | 0.003071 | -0.14659 |
| GAPDH | 22 | 0.007723 | -0.40081 |

**Table S2 Primers Used for sgRNA Expression Vector Construction. Related to Figures 1-4.**

| Guide No. | Gene Target | Strand | Primers for sgRNA cloning into pLentiGuide-Puro | SOURCE |
|---|---|---|---|---|
| 1 | SOD2 | sense | 5’-CACCGCCACCATTGAACTTCAGTGC-3’ | Eton Bioscience |
| 1 | SOD2 | antisense | 5’-AAACGCACTGAAGTTCAATGGTGGC-3’ | Eton Bioscience |
| 2 | SOD2 | sense | 5’-CACCGATGATCTGCGCGTTGATGTG-3’ | Eton Bioscience |
| 2 | SOD2 | antisense | 5’-AAACCACATCAACGCGCAGATCATC-3’ | Eton Bioscience |
| 1 | NRF2 | sense | 5'-CACCGCACATCCAGTCAGAAACCAG-3' | Eton Bioscience |
| 1 | NRF2 | antisense | 5'-AAACCTGGTTTCTGACTGGATGTGC-3' | Eton Bioscience |
| 1 | Non-targeting Control | sense | 5'-CACCGGAAAGACTATTTCAAGCAGA-3' | Eton Bioscience |
| 1 | Non-targeting Control | antisense | 5’-AAACTCTGCTTGAAATAGTCTTTCC-3' | Eton Bioscience |
